# Biocontrol of *Aspergillus niger* in 3D-lung cell tissues by oxalotrophic bacteria

**DOI:** 10.1101/2020.08.20.259929

**Authors:** Fabio Palmieri, Ilona Palmieri, Nourine Noormamode, Aislinn Estoppey, M. Omar Ishak, Julia M. Kelliher, Armelle Vallat, Rashi Iyer, Saskia Bindschedler, Karen Davenport, Patrick S. G. Chain, Jennifer Foster Harris, Pilar Junier

## Abstract

*Aspergillus* fungi are opportunistic pathogens that affect a large number of people worldwide. Many aspects of *Aspergillus* spp. pathogenesis toward humans are known, but their ability to enhance their infectious potential by manipulating the environmental pH of its host has not been considered yet. In this study, we tested the hypothesis that by producing oxalic acid, *Aspergillus niger* can manipulate pH during lung infection and thus, interfering with this process could limit pathogenicity. To test this hypothesis, we co-cultured *A. niger* with oxalotrophic bacteria in increasingly complex testing systems (Petri dishes and 3D-cell cultures systems). In *in vitro* tests, oxalotrophic bacteria limit oxalic acid production and suppressed the pH shift induced by *A. niger*. In 3D-cell cultures (Transwells® and Bronchioles-on-a-chip), *A. niger* also modified pH, Ca^2+^ and oxalic acid concentrations. Co-inoculation with as little as 10 cells of the oxalatrophic bacterium strongly inhibited the germination and development of *A. niger* and returned each of the three parameters to the baseline physiological values of uninfected cells. This biocontrol interaction between oxalotrophic bacteria and oxalate-producing *A. niger* could represent a paradigm shift in the fight against opportunistic fungal pathogens, where the host environment is rendered less permissive to fungal development.

## Introduction

Fungal diseases are estimated to kill more than 1.5 million people every year (1, 2). Over the last three decades, the increase in the number of at-risk individuals has correlated with an intensification in the burden of fungal disease on human health (3). As the size of at-risk populations (e.g. immunosuppressed patients) is expected to keep increasing in the future (2), tackling fungal pathogenesis is urgent. However, only a very limited number of antifungal drugs are used nowadays to control fungal pathogens. Because of this restricted chemical arsenal, the same classes of molecules are used in human health, animal husbandry, and agriculture, leading to the rise, and rapid spread, of resistance in fungal pathogens that can affect both plant and animal hosts (4). In spite of this, tackling fungal diseases has been largely neglected up to now (5).

The most prevalent fungal pathogens affecting humans are airborne opportunists such as *Aspergillus* spp., *Cryptococcus* spp., *Pneumocystis* spp., and human-associated commensals like *Candida albicans* (6, 7). These fungal species are responsible for approximately 90% of the deaths due to fungal infection (1). The ecology of all these organisms plays a very significant role in their ability to transition to pathogenic lifestyles. For instance, the remarkable plasticity in the ecology and stress-response of *Aspergillus* spp. is believed to form the basis of its success as an opportunistic pathogen (8, 9).

Although many aspects of the ecology of *Aspergillus* spp. have been connected to pathogenicity (9), its ability to manipulate pH via the secretion of low molecular weight organic acids (LMWOA), and in particular, oxalic acid, has been largely ignored in the context of human pathogenesis. Several clinical reports have shown the presence of calcium oxalate (CaOx) crystals in pulmonary aspergillosis in animals and humans (10-15). Oxalic acid and oxalate crystals are thought to directly cause damage to the host tissues (including pulmonary blood vessels), and to generate free radicals which can harm cells indirectly (14). A recent case report of invasive pulmonary aspergillosis in a 69-year old man with lymphoma and pneumonia indicated the presence of CaOx crystals around blood vessels and within the blood vessel walls. This suggests a potential mechanical role of oxalate crystals in the angioinvasion of *Aspergillus* (15). However, a link between oxalic acid production and pathogenicity has not yet been made in fungi from this genus or in any other fungal human pathogen. On the contrary, oxalic acid production has been widely acknowledged as a pathogenicity factor in fungal plant pathogens, such as *Sclerotinia sclerotiorum* and *Botrytis cinerea* (16, 17) where pH manipulation and calcium chelation plays a direct role in pathogenesis (18-20). Both acidification and cation complexation in the local environment is exploited by plant pathogens to weaken the cell wall structure, facilitate infection, inhibit plant defenses, and induce programmed cell death (19, 21).

Oxalic acid is a ubiquitous compound in the environment and is thought to have a central role in fungal metabolism (19). Its production and consumption by microorganisms have been directly associated with pH regulation in soil (22). In such ecosystems, oxalic acid is often found complexed with divalent cations, especially calcium (23). Despite its chemical stability (K_sp_ CaOx monohydrate = 2.32 × 10^−9^), CaOx rarely accumulates (24, 25). This is because oxalate is used by soil oxalotrophic bacteria as carbon and energy sources. The transformation of a strong organic acid into a weaker one (oxalic acid *versus* carbonic acid; pK_a_ = 1.25 and 4.14, respectively), leads to a local increase in soil pH. This overall process has been coined out in the oxalate-carbonate pathway (26). Moreover, oxalic acid plays a key role in bacterial:fungal interactions acting as a signaling cue by bacteria in order to localize fungi and to establish different types of trophic interactions with them (27-29).

In this study we wanted to evaluate whether the metabolic processes associated to oxalic acid during *Aspergillus* spp. infection could parallel those occurring during its natural cycling in the oxalate-carbonate pathway in soils. Indeed, humans can be seen as a complex ecosystem governed by the same ecological principles affecting any other ecosystem (30). Thus, the ability of oxalotrophic bacteria to degrade oxalic acid produced by *Aspergillus* spp., and control the subsequent pH manipulation and calcium chelation, would result in a mechanism to control biologically this opportunistic fungal pathogen (Fig. 1). To test our hypothesis, we selected *Aspergillus niger* as a model. This organism is regarded as a safe relative of the more infectious *Aspergillus fumigatus* (31), and has been extensively studied for its ability to produce oxalic acid (32, 33). It is also a known agent of aspergillosis in humans, being responsible for 5% of the cases (9, 34). We first confirmed that *A. niger* secretes oxalic acid and assessed the effect of co-cultivation with oxalotrophic bacteria on pH and oxalic acid concentration, and on the inhibition of *A. niger*’s growth. This was done *in vitro* and on human bronchial epithelial cell (HBEC) cultures using two complementary 3D-cell cultures systems (Transwell® inserts and bronchioles-on-a-chip (BoC)).

**Fig. 1.**
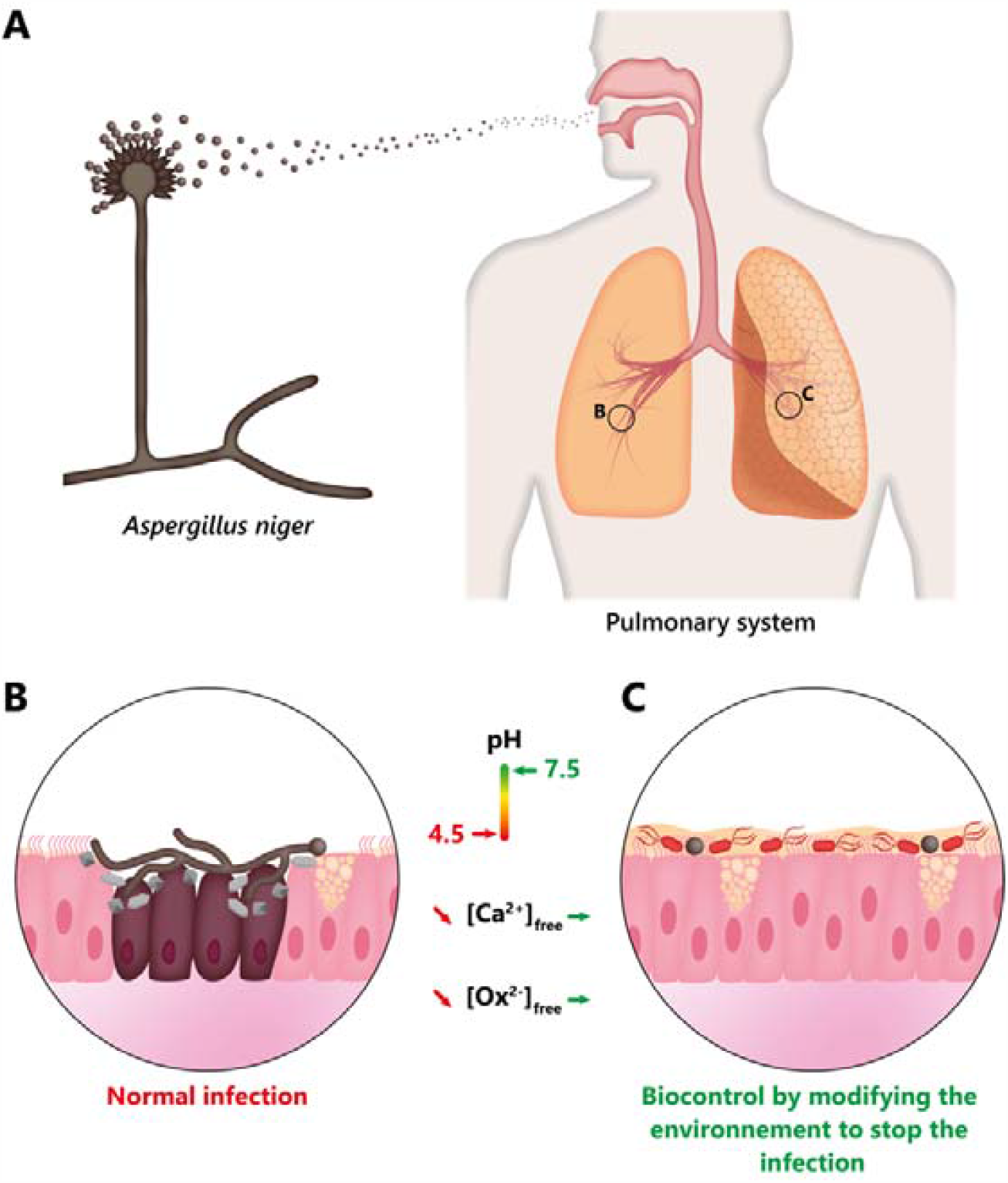
Schematic summary of the proposed strategy to control *Aspergillus niger* infection by introducing oxalotrophic bacteria to modify the *A. niger* environmental niche. (A) *A. niger* conidia (depicted in dark grey) arrive in the respiratory system through breathing. (B) During a normal infection process in a susceptible host, *A. niger* modifies the environment by secreting oxalic acid (or oxalate Ox^2+^) which decreases pH and chelates free calcium in the form of CaOx (depicted in light grey) crystals. This results in the infection of the host’s tissue. (C) The biocontrol strategy proposed here takes advantage of the ability of oxalotrophic bacteria (cells depicted in red) to consume CaOx and thus reestablish physiological pH and free calcium concentrations.

## Results

### Detection of oxalic acid produced by *Aspergillus niger* in different culture conditions

Since the metabolism of fungi can change significantly depending on the nutritional conditions of the growth medium, we first tested whether our fungal strain produced oxalic acid only or a mixture of different organic acids in culture media differing in their trophic conditions. While *A. niger* is known to produce high amounts of oxalic acid, it is also well known to produce other LMWOA such as citric acid (32, 35-37). An Ultra-High-Performance Liquid Chromatography (UHPLC) analysis revealed that in the conditions we tested, *A. niger* produced oxalic acid, only (Fig. S1A). The absence of other organic acids is not surprising as the conditions for other LMWOA production are highly specific (e.g. carbon concentration above 50 g/L carbon and pH < 3; (38, 39) and are not provided in the conditions we tested. Acidification and the presence of CaOx crystals could also be detected in water yeast agar (WYA) (Fig. S1B). We thus concluded that our *A. niger* strain consistently produced oxalic acid and acidified the pH of its medium under laboratory growth conditions.

### Confrontation assays between *A. niger* and selected bacterial strains

We assessed how the presence of oxalotrophic bacteria impacted *A. niger* growth and pH evolution in different growth media. Two oxalotrophic bacteria (*Cupriavidus necator* and *C. oxalaticus*) and one non-oxalotrophic bacterium (*Pseudomonas putida*) were used in confrontation assays. In malt agar diluted 10 times (MA 1/10), *A. niger* colonized the entire Petri dish, including the area in which the bacterial inocula were applied. All three inoculated bacteria did not survive in the area of interaction with the fungus (Fig. S2A). In Reasoner’s 2 agar (R2A), *A. niger* mycelia did not colonize the area beyond the barrier formed by the bacterial inocula, but some hyphae were still able to grow beyond the bacterial inoculation zone and develop into microcolonies in the co-culture with *P. putida* (Fig. S2B). In WYA, mycelial growth was also restricted to the area delimited by the inocula in the co-culture with the two oxalotrophic bacteria. On the other hand, *P. putida* did not survive the interaction with the fungus, which instead, colonized the entire plate. Moreover, in WYA medium, which contained a pH indicator, the acidification of the medium by the fungus grown alone or in co-culture with the non-oxalotrophic bacterium was clearly visible. In contrast, in the co-cultures with oxalotrophic bacteria, the pH of the medium did not change. The control over fungal growth was particularly remarkable in the case of *C. oxalaticus* (Fig. S2C). Given the impact of the medium on the control of fungal growth, we repeated the confrontation assays in Air-Liquid Interface (ALI) medium, which is the medium used for differentiation of lung cells (40). The non-oxalotrophic bacterial model (*P. putida*) acidified the medium when grown alone, while the oxalotrophic bacteria (*C. necator* and *C. oxalaticus*) did not acidify the medium (Fig S3A). *C. oxalaticus* was found to not only control mycelial growth but also inhibit conidia germination (Fig. S3B). Moreover, the presence of *C. oxalaticus* in co-culture with the fungus stabilized the pH of the culture medium at a neutral pH (Fig. S3C), consistent with decreased oxalic acid concentration in the medium (Fig. S3D). We therefore conclude that oxalotrophic bacteria have a significant inhibitory effect on *A. niger* growth in all media tested.

### Establishment of a dose-response curve on submerged cultures

In order to obtain evidence that oxalotrophic bacteria control the growth of *A. niger* in an animal-free lung infection model that is compatible with the 3R principles of animal experimentation (41), we set up experiments on human bronchial epithelial cell (HBEC) cultures. We established a dose-response curve for increasing conidial and/or bacterial loads on HBECs in submerged cultures, to identify the optimal load to perform experiments in Transwells® and BoC systems. After 24 h, the overall size, shape and integrity of the lung cells changed at an absolute load of 500 conidia and above. The HBECs shrank in size due to actin agglomeration. Moreover, from a conidial load of >=1000, fungal growth also had an adverse effect on tissue integrity (Fig. S4). The same experiment was performed with *C. oxalaticus* and *P. putida*. A strong morphological change was induced by a total load of 500 bacterial cells and above for the former (Fig. S5), and as little as 10 cells for the latter (Fig. S6), which was not used further. The HBECs co-cultured with *C. oxalaticus* became rounder, and actin agglomeration increased compared with the cells-only control (Fig. S5).

To analyze the effect of co-culturing *A. niger* with the oxalotrophic bacterium on HBECs integrity, we performed a test with 10 and 500 conidia confronted with 10 bacterial cells. After 72h, *A. niger* induced morphological changes (size reduction and actin agglomeration), with a stronger effect for 500 conidia, confirming the results obtained at 24h. With the co-inoculation of as few as 10 *C. oxalaticus* cells, the morphology of the HBECs was similar to the morphology of HBECs of the bacteria-only control, suggesting the inhibition of fungal development (Fig. S7). We concluded that a conidial and bacterial load of 10 conidia/cells was ideal to monitor the interaction of *A. niger* and *C. oxalaticus* in differentiated HBECs in Transwells® and BoC systems.

### Biocontrol assay of *A. niger* bronchial cells infection by *C. oxalaticus*

After establishing a dose-response curve on HBECs in submerged cultures, the effect of inoculation of 10 *A. niger* conidia alone or in co-culture with 10 *C. oxalaticus* cells was assessed in differentiated bronchial tissue in Transwell® inserts and BoC systems. In the presence of the fungus alone, changes in three key environmental factors were observed: pH, Ca^2+^ concentration, and concentration of soluble oxalic acid. The pH dropped significantly from 7.5 down to 4.5 in Transwells® and from 7.3 to 6.8 in BoC systems (Fig. 3A). Ca^2+^ concentrations changed from 1 mM to around 0.2 mM in both culture systems (Fig. 3B). The level of soluble oxalic acid produced by *A. niger* dropped from 500 μM (cells alone) to 75 μM (cells inoculated with the fungus) (Fig. 3C). In contrast, pH, Ca^2+^ and free oxalic acid levels were statistically indistinguishable when oxalotrophic bacteria were co-cultured with the fungus compared to the controls with lung cells alone or with bacteria. In addition, CaOx crystals were observed in the cultures in which the fungus developed (Fig. 3D), but not when the fungus was co-cultured with oxalotrophic bacteria (Fig. 3E). This suggests that the lower levels of soluble oxalic acid measured in the treatment with the fungus were likely the result of complexation of oxalic acid and Ca^2+^, and corroborates the pH and Ca^2+^ concentration data. Moreover, the absence of CaOx crystals when the fungus was in co-culture with *C. oxalaticus* agrees with the pH, Ca^2+^ concentration and soluble oxalic concentrations measured. These results validated our hypothesis that oxalotrophic bacteria can be used to manipulate the microenvironment created by *A. niger*. In addition to the changes in the environmental parameters measured above, we also observed a cytopathic effect when conidia of *A. niger* developed into mycelia (Fig. 4A and B). This cytopathic effect resulted in the destruction of the bronchial epithelium. Lactate dehydrogenase (LDH) activity were elevated in response to the presence of the foreign oxalotrophic bacteria (Fig. 4C), something that needs to be addressed for any future therapeutic application.

**Fig. 2.**
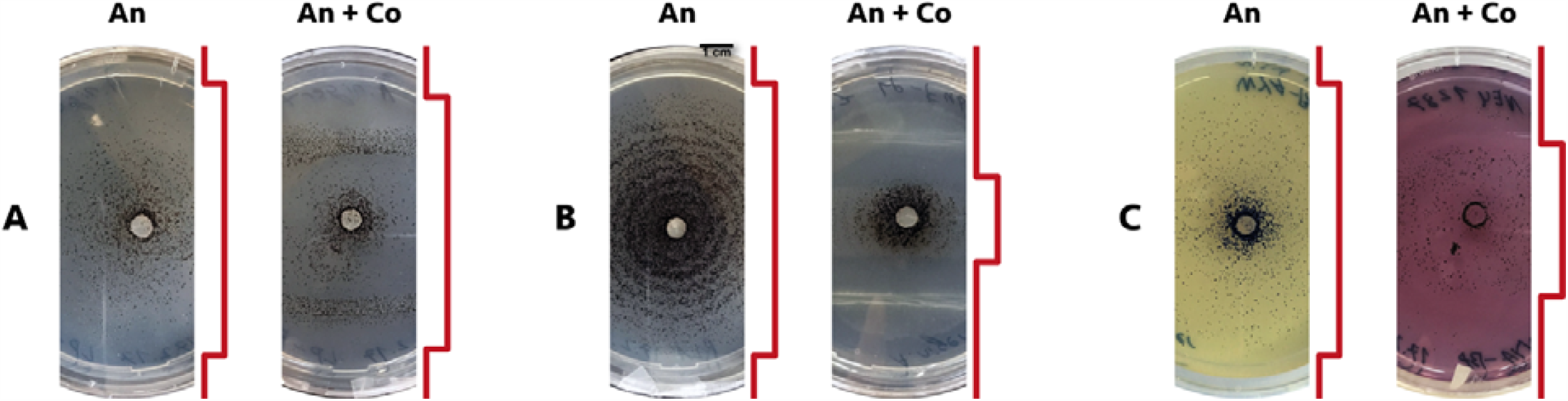
Comparison of the growth of *Aspergillus niger* (An) alone and in confrontation with *Cupriavidus oxalaticus* (Co) in different culture media. The red line next to each picture represents the extent of *A. niger* growth. On MA 1/10 (A), there is no significant growth inhibition of *A. niger*, as it kills *C. oxalaticus. A. niger* growth is highly restricted to the center of the plate when co-cultured with *C. oxalaticus* on R2A (B). The growth inhibition of *A. niger* when co-cultured with *C. oxalaticus* is less pronounced on WYA + BP. Moreover, the presence of C. oxalaticus revert the pH of the medium to a neutral value (C). A yellow color indicates an acidic pH <6.

**Fig. 3.**
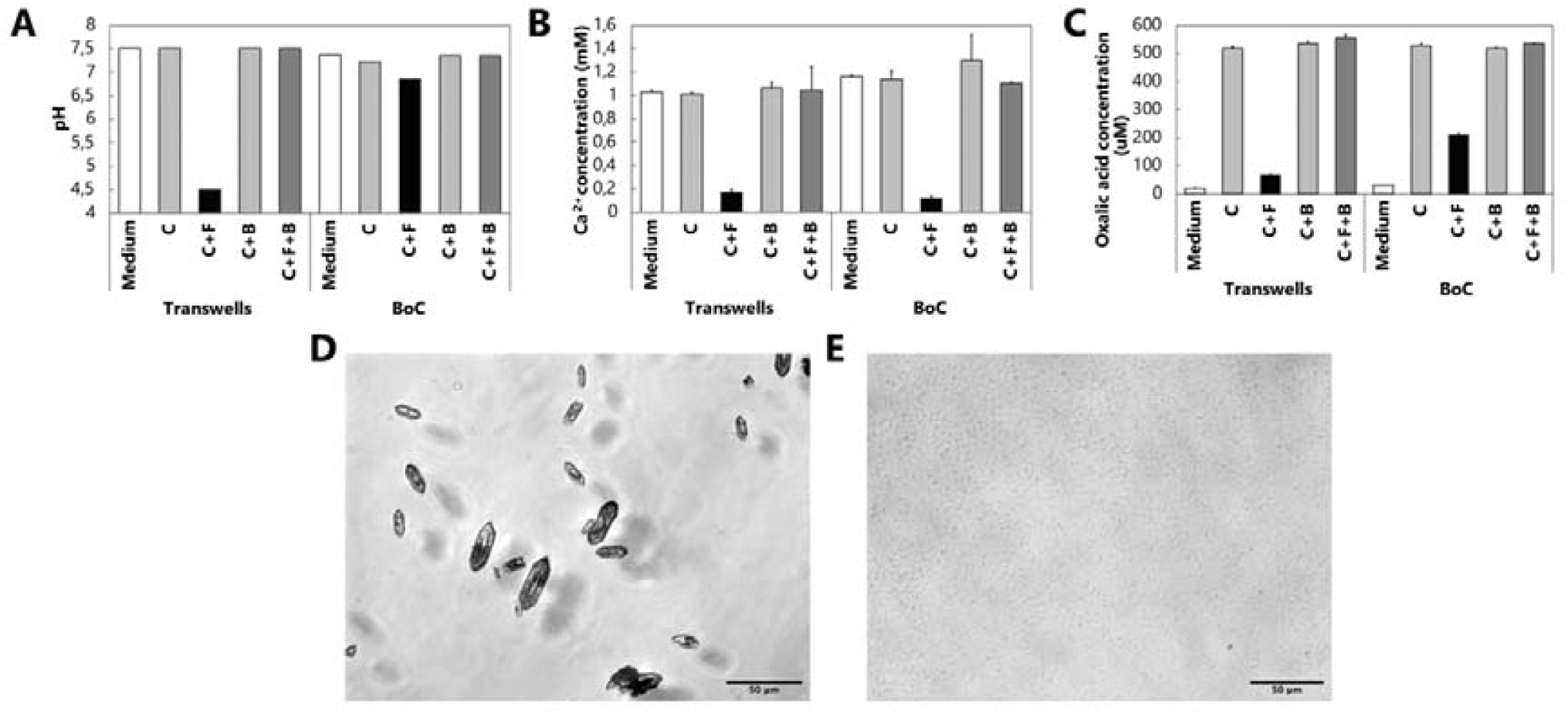
Influence of the interaction between *Aspergillus niger* and the oxalotrophic bacterium *Cupriavidus oxalaticus* on environmental parameters of differentiated bronchial tissue in Transwell® inserts and bronchiole-on-a-chip (BoC) devices. In the presence of the fungus, the pH (A) decreases, as compared to all other treatments. This pH decrease is correlated with a drastic decrease in the concentration of free Ca^2+^ (B) (*p*-value between C and C+F = 8,951 × 10^−7^). Free oxalic acid concentrations were lower in the presence of the fungus, compared with the basal level secreted by the bronchial cells (C) (*p*-value between C and C+F = 4,587 × 10^−8^). These results are supported by the detection of CaOx crystals in the presence of the fungus (D). In the co-culture with the oxalotrophic bacterium, pH, free Ca^2+^ and free oxalic acid concentrations return to physiological levels, and this was concomitant with the absence of crystals (E) (*p*-values between C+F and C+F+B for free Ca^2+^ and free oxalic acid = 0,002 and 1,415 × 10^−7^, respectively). C: lung cells; C+F: lung cells + fungus; C+B: lung cells + bacteria; C+F+B: lung cells + fungus + bacteria. For A, B, C, the results represent the mean + sd of three independent measurements for the Transwell® inserts for each condition (three biological replicates, n = 3). For A, pH results for the BoC devices represent a unique measurement per condition (one replicate, n = 1). For B and C, Ca^2+^ and oxalic acid results for the BoC devices represents the mean + sd of two measurements per condition (one replicate).

**Fig. 4.**
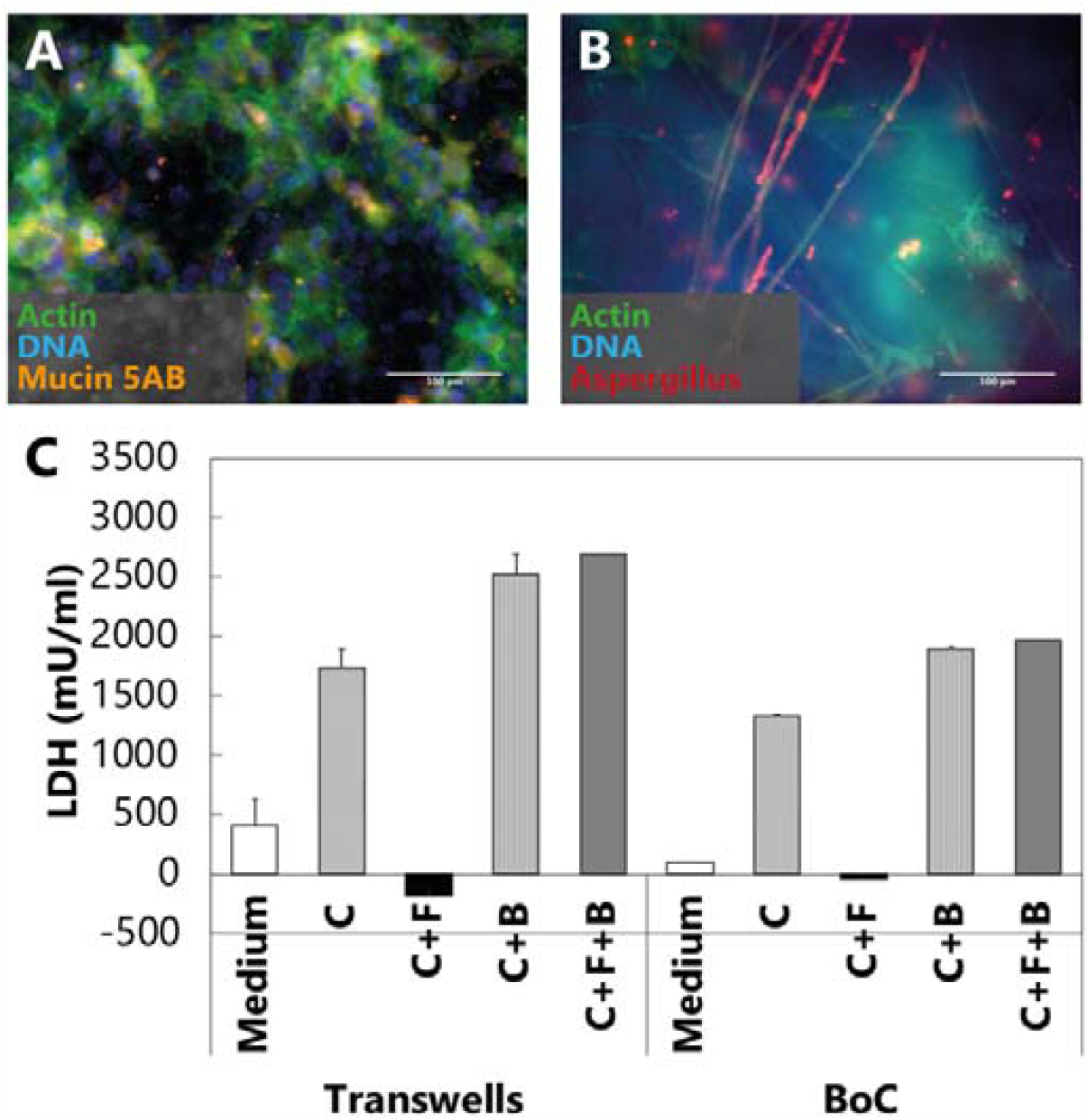
Cytopathic effect of *Aspergillus niger* on differentiated bronchial tissue and lactate dehydrogenase (LDH) measurements of the co-culture between *A. niger* and *Cupriavidus oxalaticus* in Transwell® inserts and bronchiole-on-a-chip (BoC) devices. (A) Control showing healthy differentiated bronchial epithelial cells. (B) Bronchial epithelial cells infected with *A. niger*. (C) LDH leakage was measured as a proxy for cell damage. In the presence of the fungus, no LDH has been detected, probably because of the destruction of the tissue by *A. niger* (B). *C. oxalaticus* cause significantly more LDH leakage that the basal LDH level of control cells (p-value = 0,004). C: lung cells; C+F: lung cells + fungus; C+B: lung cells + bacteria; C+F+B: lung cells + fungus + bacteria. For C, the results represent the mean + sd of three independent measurements for the Transwell® inserts for each condition (three biological replicates, n = 3). LDH results for the BoC devices represents the mean + sd of three measurements per condition (one replicate, n = 1).

### Genomic potential for oxalic acid production in other *Aspergillus* spp

We performed a genomic screening of orthologous genes to the oxaloacetate acetylhydrolase (*oahA*) and the oxalate/formate antiporter (genes involved in oxalic acid production in *A. niger*) in genomes available in the *Aspergillus* Genome Database (AspGD). The genomic screening revealed that orthologs of both genes are found in multiple *Aspergillus* spp. and are highly conserved across diverse species (Fig. 5). In the case of the orthologs to the oxalate/formate antiporter, they were conserved to a lesser extent (Fig. S8). This genomic analysis confirmed that diverse *Aspergillus* spp. possesses the genes necessary to produce and secrete oxalic acid.

**Fig. 5.**
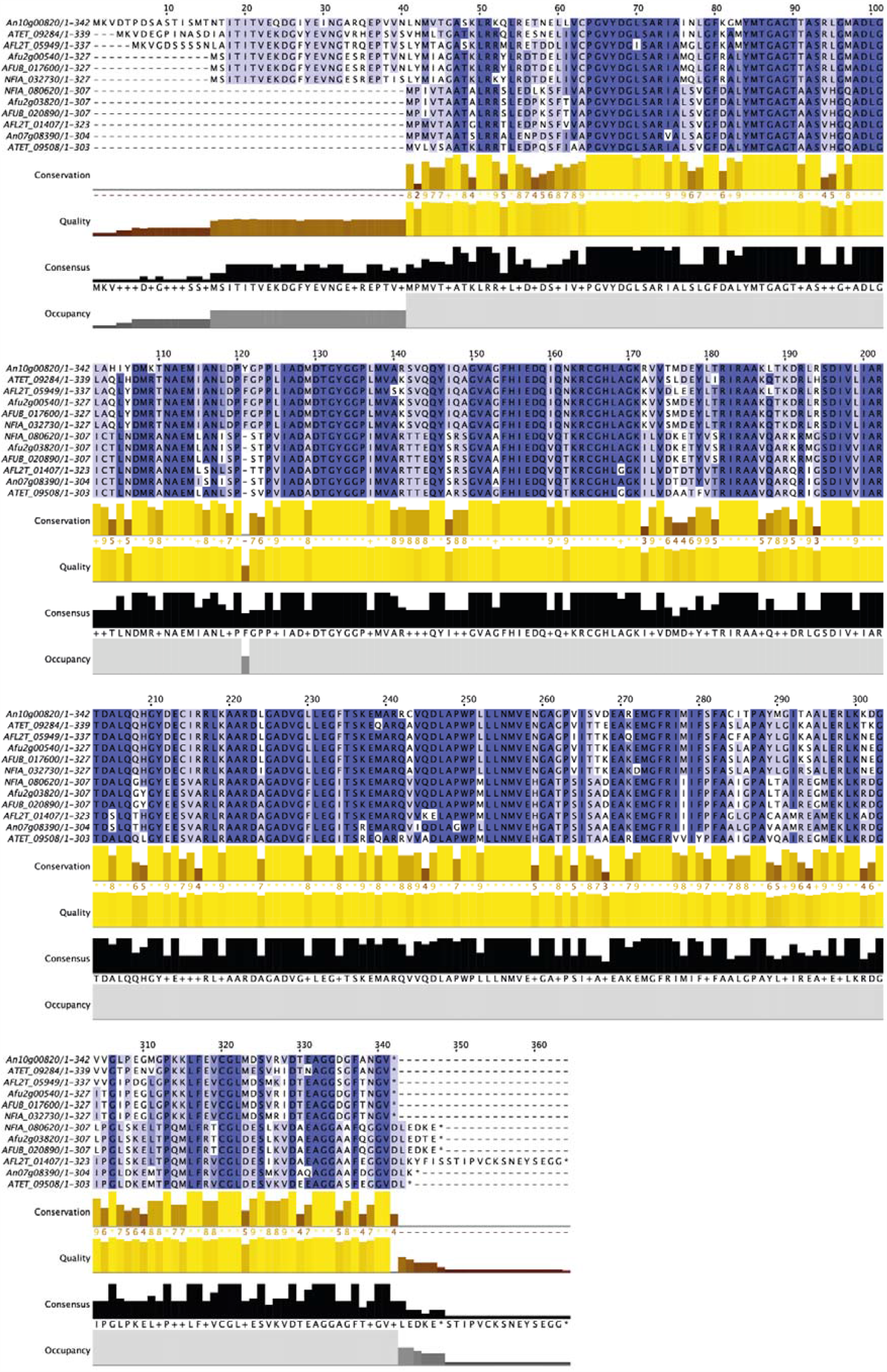
Genomic screening of the oxaloacetate acetylhydrolase (OAH) in other *Aspergillus* spp. Multiple sequence alignment of the protein sequences orthologous to the OAH of *A. niger* CBS 513.88 (GenBank accession number CAD99195.1) revealed they were well conserved across diverse species, as indicated by an intense purple color of the amino acids. Multiple sequence alignments were performed using the MUSCLE protein alignment algorithm in Jalview (version 2.11.1.2). An = *A. niger* CBS 513.88, ATET = *A. terreus* NIH2624, AFL2T = *A. flavus* NRRL 3357, Afu = *A. fumigatus* Af293, AFUB = *A. fumigatus* A1163, NFIA = *Neosartorya fisheri* NRRL 181 (formerly *A. fisheri*).

## Discussion

Here we present a biological interaction between *A. niger* and oxalotrophic bacteria that results in the biological control of *A. niger*, preventing infection in 3D-lung cell tissues (Transwells® and BoC). The direct consequence of acidification through oxalic acid production by *A. niger* was the decrease in free Ca^2+^ and subsequent precipitation of CaOx crystals. CaOx crystals are well known to occur in lung tissues upon infection by *A. niger* (10, 11, 13). Presumably, by consuming the oxalate produced by *A. niger*, the oxalotrophic bacterial species *C. oxalaticus* blocks the subsequent decrease in pH and formation of CaOx crystals observed in the absence of the bacterium. To obtain a direct confirmation of the role of oxalic acid in the manipulation of pH during lung infection, the use of a non-oxalate-producer *A. niger* mutant would be indispensable. Such mutants (*oahA* gene) are described in the literature and exhibited a decreased acidification of the culture medium and reduced extracellular protease activity (32, 42). After multiple failed attempts to obtain the published mutants by addressing the corresponding scientific teams, we attempted to construct a non-oxalate-producer mutant of our *A. niger* strain using CRISPR-Cas9 gene editing. However, this was unsuccessful due to multiple targets of the sgRNA probes and thus could not be included in this study.

Oxalate-degrading bacteria are known inhabitants of the human gut, where they perform the key function of degrading dietary oxalate (43). These species have also been used as probiotics for the treatment of hyperoxaluria (high oxalate in urine) and the management of kidney stones (43, 44). While oxalate-degrading bacteria are well characterized in the gut, this is not the case of the lung. Although considered sterile for a long time, the lung is now known to harbor a diverse microbiota (45, 46). Oxalate-degrading capabilities have been previously reported in strains of the genera *Lactobacillus* (47), *Streptococcus* (48), *Prevotella* (49, 50) and *Veillonella* (50), all of which are reported as components of the lung microbiota. However, assessing the oxalotrophic potential of the lung microbiome is something that still need to be accomplished.

The genomic analysis of multiple *Aspergillus* spp. suggests that oxalotrophy could also be relevant to other *Aspergillus* causing pulmonary aspergillosis (5). The presence of CaOx crystals during infection by *A. fumigatus* has been reported in the literature (8, 11, 14, 51). Accordingly, we found orthologs of the oxaloacetate acetylhydrolase (OAH) and the oxalate/formate antiporter of *A. niger* in the genomes of two well characterized model *A. fumigatus* strains Af293 and A1163 (52), suggesting the production of oxalic acid by this pathogen and the potential of using oxalotrophic bacteria in fungal species more relevant for human health. To conclude, the results presented here represent a stepping stone towards developing an alternative approach to control the development of oxalate-producing *Aspergillus* spp. based on the manipulation of the lung environment using bacterial:fungal interactions.

## Materials and Methods

### Bacterial and Fungal cultures

All bacterial and fungal strains come from the collection of the Laboratory of Microbiology of the University of Neuchâtel (LAMUN; Table 1). *P. putida* KT2440 was kindly provided by Dr. Arnaud Dechesne (Technical University of Denmark). *C. necator* JMP289 was kindly provided by Prof. Jan van der Meer (University of Lausanne). *C. oxalaticus* Ox1 was tagged in-house using insertion with a MiniTn7 system. Table 2 summarizes all the media used. Bacterial strains were routinely cultured on NA medium. *Aspergillus niger* was routinely cultured on MA medium. PDA was used for *A. niger* conidia production. BHIA was used to have mycelium-only colony edge without any conidia in order to prevent unwanted conidia dispersal during confrontations with bacteria.

**Table 1.**
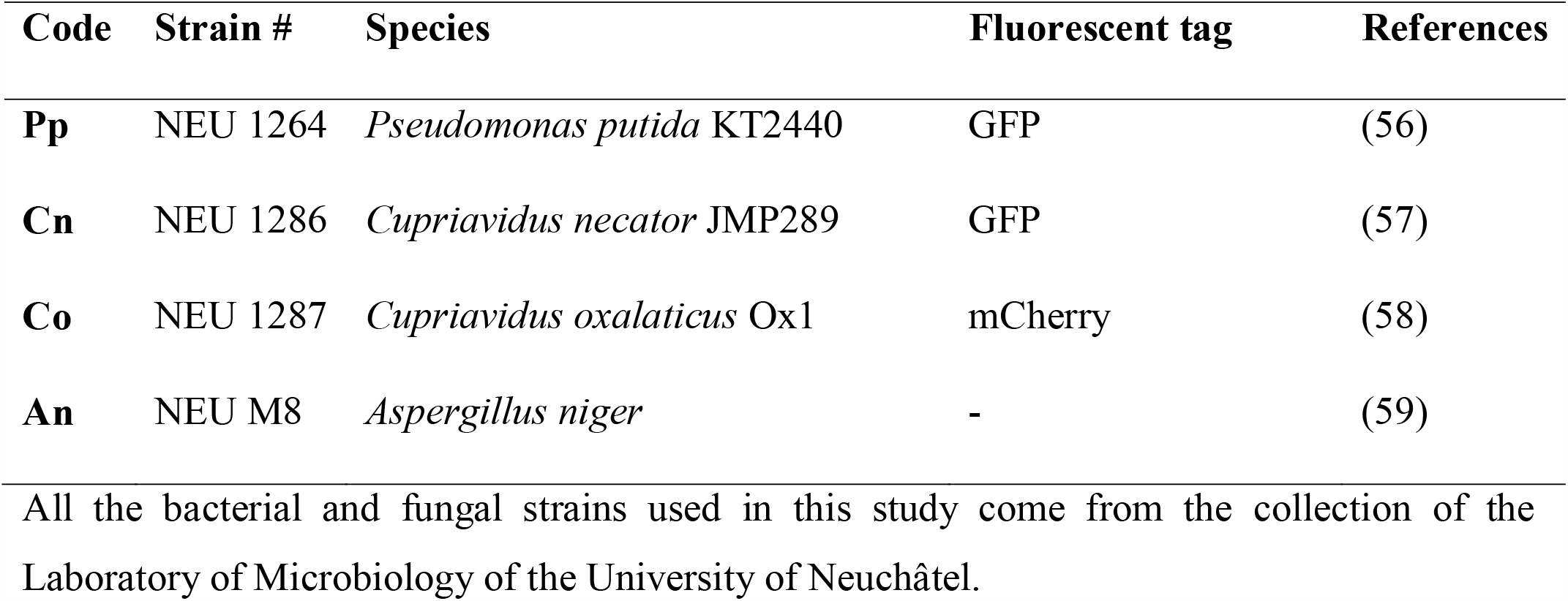
Bacterial and fungal strains used.

| Code | Strain # | Species | Fluorescent tag | References |
| --- | --- | --- | --- | --- |
| **Pp** | NEU 1264 | *Pseudomonas putida* KT2440 | GFP | (56) |
| **Cn** | NEU 1286 | *Cupriavidus necator* JMP289 | GFP | (57) |
| **Co** | NEU 1287 | *Cupriavidus oxalaticus* Ox1 | mCherry | (58) |
| **An** | NEU M8 | *Aspergillus niger* | - | (59) |

All the bacterial and fungal strains used in this study come from the collection of the Laboratory of Microbiology of the University of Neuchâtel.

**Table 2.**
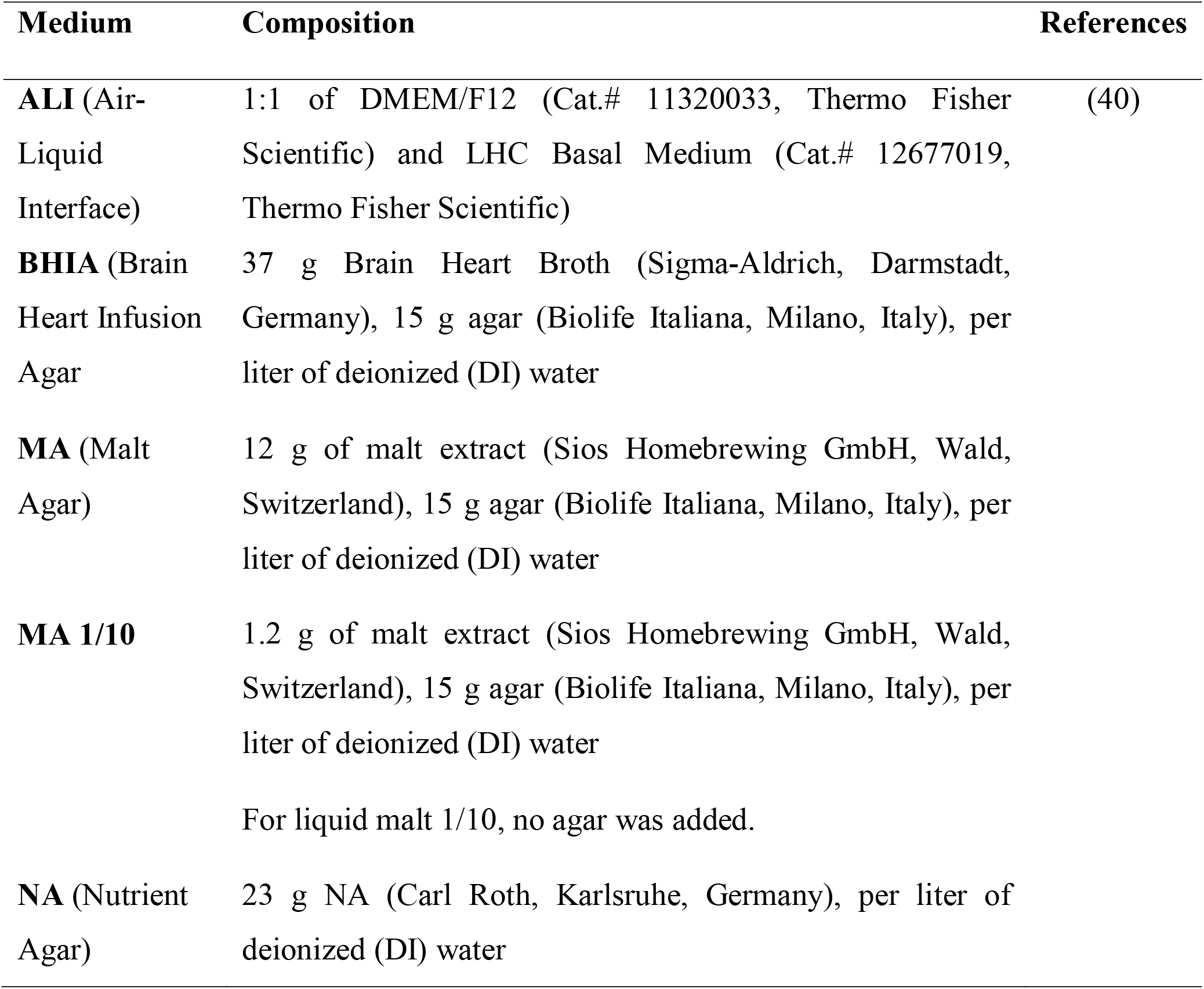
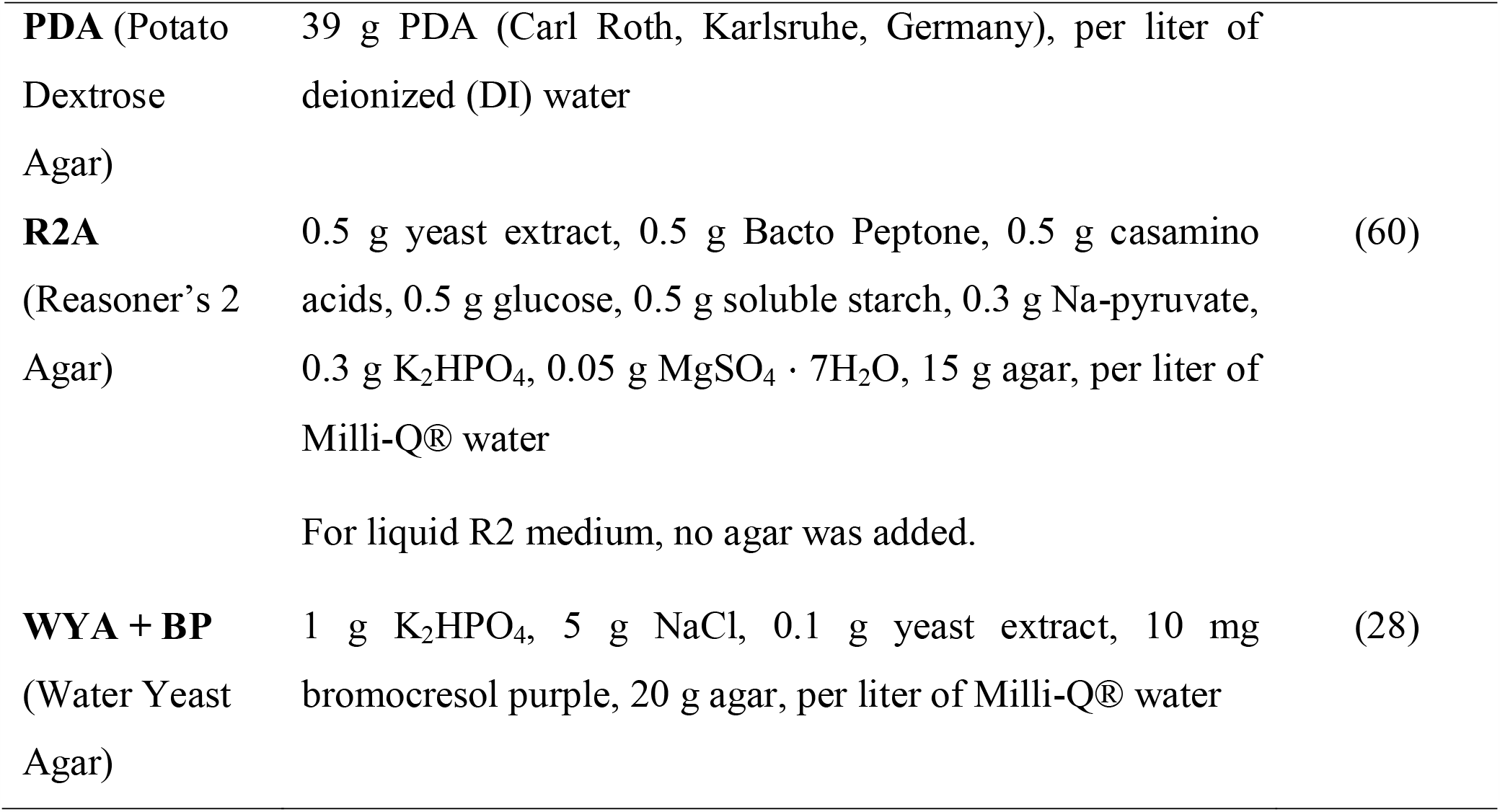
Culture media recipes.

| Medium | Composition | References |
| --- | --- | --- |
| **ALI** (Air-Liquid Interface) | 1:1 of DMEM/F12 (Cat.# 11320033, Thermo Fisher Scientific) and LHC Basal Medium (Cat.# 12677019, Thermo Fisher Scientific) | (40) |
| **BHIA** (Brain Heart Infusion Agar) | 37 g Brain Heart Broth (Sigma-Aldrich, Darmstadt, Germany), 15 g agar (Biolife Italiana, Milano, Italy), per liter of deionized (DI) water |  |
| **MA** (Malt Agar) | 12 g of malt extract (Sios Homebrewing GmbH, Wald, Switzerland), 15 g agar (Biolife Italiana, Milano, Italy), per liter of deionized (DI) water |  |
| **MA 1/10** | 1.2 g of malt extract (Sios Homebrewing GmbH, Wald, Switzerland), 15 g agar (Biolife Italiana, Milano, Italy), per liter of deionized (DI) water<br>For liquid malt 1/10, no agar was added. |  |
| **NA** (Nutrient Agar) | 23 g NA (Carl Roth, Karlsruhe, Germany), per liter of deionized (DI) water |  |
| **PDA** (Potato Dextrose Agar) | 39 g PDA (Carl Roth, Karlsruhe, Germany), per liter of deionized (DI) water |  |
| **R2A** (Reasoner's 2 Agar) | 0.5 g yeast extract, 0.5 g Bacto Peptone, 0.5 g casamino acids, 0.5 g glucose, 0.5 g soluble starch, 0.3 g Na-pyruvate, 0.3 g $K_2HPO_4$, 0.05 g $MgSO_4 \cdot 7H_2O$, 15 g agar, per liter of Milli-Q® water | (60) |
|  | For liquid R2 medium, no agar was added. |  |
| **WYA + BP** (Water Yeast Agar) | 1 g $K_2HPO_4$, 5 g NaCl, 0.1 g yeast extract, 10 mg bromocresol purple, 20 g agar, per liter of Milli-Q® water | (28) |

### LMWOA detection by UHPLC and by colorimetric pH indicator-based Petri dish assay

For the UHPLC analysis, 500 µl of 30 mM H_2_SO_4_ were added to 1 mL of a two-week liquid culture in malt 1/10, Reasoner’s 2, and ALI liquid media in triplicate, to obtain 20 mM H_2_SO_4_ final concentration in order to obtain a low pH for the extraction of LMWOAs and to dissolve any precipitated crystals. The samples were incubated at 60°C for two hours to dissolve precipitated metal oxalate crystals, and then centrifuged at 3000 g for 10 min. All the samples were filtered at 0.22 µm (13mm syringe filters, PTFE, hydrophilic) and 200 µl were added into HPLC vials with 250 µl conical inserts. UHPLC (Ultimate 3000 RS-Dionex, Thermo Fisher Scientific, USA) was coupled with DAD detector set at 210 ± 2 nm. A 5 µL of sample was injected onto a Prevail− organic acid column (5 µm particle size, 150 x 4.6 mm, Grace Davison Discovery Sciences, USA) with the temperature kept at 40°C. The mobile phase consisted of 50 mM phosphate buffer adjusted to pH 2.5 with phosphoric acid with a flow rate of 1 mL/min. Pure oxalic acid (Merck, Germany) was identified by the retention time and was quantified by an external standard curve, linear regression from five calibration points (0.2 to 5 mg/mL). For the culture-based assay, WYA supplemented with bromocresol purple (WYA+BP) was used as a pH indicator-containing medium. After one week of incubation at room temperature (RT), the presence of typical bi-pyramidal shaped CaOx crystals was assessed by observing a thin slice of agar medium sampled at the edge of the colony and stained with lactophenol cotton blue under a Leica DM4 B optical microscope connected to a Leica DFC7000 T camera.

### Confrontation assays on solid media

Confrontations assays were performed between *A. niger* and *P. putida, C. necator* and *C. oxalaticus* (Table 1), on three culture media (MA 1/10, R2A and WYA+BP). A plug coming from the apical part of an actively growing *A. niger* colony was sampled using the wider end of a Pasteur pipette and inoculated in the center of the plates. The bacterial strains were inoculated from fresh plates as opposite lines on either side of the fungal inoculum. Plates were incubated at RT for 20 days, and pictures of the plates at 20 days were taken. Pictures of the bacterial inocula were taken using a Nikon SMZ18 epifluorescence stereoscope, connected to a Nikon DS-Ri2 camera, in order to assess the viability of the bacterial strains thanks to constitutively expressed fluorescent proteins.

### Growth tests and confrontation assay in Air-Liquid Interface (ALI) medium

Bacterial growth in ALI medium was tested for 3 days at RT. To produce conidial suspensions, *A. niger* was cultured on PDA for 10 days at RT. Conidia were harvested using Dulbecco’s Phosphate Buffer Saline (DPBS) supplemented with 0.01% (v/v) Tween 80. Harvested conidia were washed three times with DPBS following centrifugation at 2000xg for 5 min at RT. Finally, conidia were resuspended in 2 mL DPBS and quantified with an Improved Neubauer counting chamber. Two μL of 16’000 conidia/bacterial cell per μL suspensions were inoculated in 200 μL ALI medium. To test mycelial growth in the different media, small agar plugs (approximately 3×3 mm) coming from the edge of a colony of *A. niger* on BHIA were used. The 96-well plate was incubated at RT for 7 days and growth was visually assessed.

Confrontation of *A. niger* with *C. oxalaticus* was performed in 100-mm Corning® tissue culture plate containing 12 mL ALI medium to allow the fungus to attach during growth. The fungal inoculum was taken from the apical part of an active colony grown on BHIA by using the wider end of a Pasteur pipette. For the confrontation assay, the fungal inoculum was placed in the medium after addition and mixing of 100 μL inoculum of an overnight culture of *C. oxalaticus* in ALI medium. A fungus-only plate was used as control. The plates were incubated at RT for 7 days. Oxalic acid concentration was quantified by using the Oxalic Acid Colorimetric Assay Kit (Sigma-Aldrich, Germany), following the manufacturer instructions.

### Preparation and sterilization of the bronchiole-on-a-chip (BoC)

The chip was assembled as described in Hsieh et al. (53). Each unit of the culture platform was fabricated by using a layer-by-layer stacking technique (54). The devices were designed using Solid Edge 2D software (ST9, Siemens PLM Software), and each layer of the prelaminated polymeric sheet was obtained using a CO_2_ laser cutter (Universal Laser System). The prelaminated polymeric sheets were combined with biocompatible adhesive tapes (9122, 3M Company) with PMMA (1.5 and 3 mm thick) or PET (0.1 and 0.25mm thick). After cutting, each layer was aligned and assembled using a seam roller to complete the devices. The culture chip includes a Y-shaped apical and a basal part separated by a porous PET membrane (pore size = 0.4μm) prepared as described in Arefin *et al*. (55). The PET membrane was sandwiched between two PET sheets using adhesive transfer tape to create the cell culture surface. This allows nutrients to pass from the media to the cells through the porous membrane. The open design of the tissue chip makes cell seeding procedure easy and accessible.

For sterilization, each chip was placed in a 100 mm Petri dish and sterilized with 5% H_2_O_2_ solution for 1 h. The chips were then rinsed 2-3 times with sterile deionized water for 15 min between each rinse. Once all liquid was removed, the chips were let dry overnight under a laminar flow hood. The next day, the inlet and outlet of the chip were connected with a sterile tubing and rinsed 3 more times with sterile deionized water as explained before. After the last rinse, 200 μl sterile DPBS was added in the channel (apical part) and 5 mL in the basolateral part (bottom part) of the chip, and the chip was placed in a humidified incubator at 37°C with 5% CO_2_ overnight. The next morning, peroxide contamination was checked in each chip using a CG8+ i-STAT cartridge (Abbott, USA). Rinses with sterile deionized water and overnight incubation with sterile DPBS were repeated until peroxide was no longer detected.

### Primary normal human bronchial epithelial cell culture

Primary normal human bronchial epithelial cells (Lifeline Cell Technology, USA) were expanded in a T-75 cell culture flask with vent cap (Corning, USA) in BronchiaLife™ B/T complete medium (Lifeline Cell Technology, USA) supplemented with 0.5% Phenol Red solution (Sigma-Aldrich, USA, 15 mg/L final concentration) to 70-80% confluence in a humidified incubator at 37°C with 5% CO_2_. Culture medium was changed every other day. Cells were used until passage 2 for all experiments. Cells were harvested by trypsinization with 0.05% Trypsin / 0.02% EDTA (Lifeline Cell Technology, USA), followed by the addition of Trypsin Neutralizing buffer (Lifeline Cell Technology, USA), and counted using a hemocytometer after centrifugation at 100xg for 5 min and resuspension of the cell pellet in BronchiaLife™ medium.

Cells were seeded at a density of 3×10^4^ cells/well in 200 μl BronchiaLife™ medium for submerged undifferentiated tissue culture in 96-well plates, and 5×10^4^ cells and 8.6×10^4^ cells in 200 μL BronchiaLife™ medium for air-lifted differentiated tissue culture in the apical side of Transwell® inserts in 24-well plates (Corning, USA) and BoC devices, respectively. Transwell® inserts and BoC were first coated with collagen (30 μg/mL) prior seeding of the cells in order to allow proper cell attachment onto the porous membrane, as described in Arefin *et al*. (55). 600 μL of BronchiaLife™ medium was added to the basolateral side of Transwell® inserts in the 24-well plates and 3 mL in the basolateral side of the BoC device. 96-well plates, Transwell® inserts, and BoC devices were placed in a humidified incubator at 37°C with 5% CO_2_ for 2-3 days until confluence and formation of a monolayer of bronchial cells. For differentiated bronchial cell tissues (Transwells® and BoCs), cells were shifted to air-liquid interface by removing carefully the BronchiaLife™ medium from the apical side and replacing it by Air-Liquid Interface (ALI) Epithelial Differentiation Medium (Lifeline Cell Technology, USA) supplemented with 0.5% Phenol Red solution (Sigma-Aldrich, USA, 15 mg/L final concentration). The same was done for the medium on the basolateral side. Finally, the medium on the apical side was removed and the inserts and devices were placed in a humidified incubator at 37°C with 5% CO_2_ for 21 days. Medium was changed every other day as described previously. The cultures were observed daily using an EVOS™ XL Core bright field inverted microscope (Thermo Fisher Scientific, USA).

### Determination of conidial and bacterial load and confrontation assay on submerged undifferentiated bronchial epithelial cell cultures

In order to determine the optimal conidial and bacterial load to be used for confrontation on bronchial tissue cultures, increasing conidial and bacterial loads were tested to assess their effect on the morphology of bronchial epithelial cells in submerged cultures. *A. niger* was cultured on PDA for 7 days at 37°C in order to produce conidia. *A. niger* conidia were then harvested as already described. Finally, conidia were resuspended in 2 mL DPBS and quantified with an Improved Neubauer counting chamber. Bacteria were cultured in BronchiaLife™ medium at 37°C overnight and quantified with an Improved Neubauer counting chamber. A stock suspension of *A. niger* conidia was made at 10^6^ conidia/mL that was diluted further to obtain suspensions at 5×10^5^ to 5×10^3^ and 10^3^ conidia/mL. The same was done for *C. oxalaticus* from a stock suspension at 10^5^ bacterial cells/mL diluted until 10^3^ bacterial cells/mL. 10 μL of each suspension was added to submerged undifferentiated bronchial tissue in a 96-well plate in order to have 10^4^ to 10 conidia/well (200 μL) for *A. niger*, and 10^3^ to 10 bacterial cells/well (200 μL) for *C. oxalaticus*. The plates were placed in a humidified incubator at 37°C with 5% CO_2_ for 24h. For the confrontations assay, cells were infected with 10 or 500 *A. niger* conidia and were put in confrontation with 10 bacterial cells. The plate was placed in a humidified incubator at 37°C with 5% CO_2_ for 72h. After incubation, cells were fixed, stained (actin and nucleus), and imaged as described below (Immunofluorescence staining).

### Confrontations on differentiated bronchial tissues in Transwell® inserts and BoC devices

Differentiated bronchial tissues in Transwell® inserts and in BoC devices were infected with 10 μL of 10^3^ conidia or bacterial cell to get 10 conidia or bacterial cells. Fungus and bacteria were co-inoculated for the confrontation and controls with only medium, cells, fungus or bacteria were included. Each condition was done in triplicate for the Transwell® inserts and one unique replicate for each condition was done for the BoC devices. All Transwell® inserts and BoC devices were incubated in a humidified incubator at 37°C with 5% CO_2_ for 72h.

### Immunofluorescence staining

Undifferentiated and differentiated bronchial tissues (Transwell® and BoCs) were fixed with 100 μL 4% paraformaldehyde in DPBS for 15 min at RT. Cells were then rinsed 3 times with 200 μL DPBS, with 2 min waiting time between each rinse. Cells were permeabilized with 100 μL 0.5% Triton X-100 in DPBS for 15 min at RT and rinsed 3 times with 200 μL DPBS, with 2 min waiting time between each rinse. After that, cells were blocked with 100 μL 3% BSA in DPBS for 1h at RT. Anti-Mucin 5AC mouse monoclonal antibody (Abcam, USA, Cat.# ab218466) was prepared in DPBS (1/100). The actin stain (ActinGreen™ 488 ReadyProbes™ Reagent) and the nuclei counterstain (NucBlue™ Live ReadyProbes™ Reagent) were added to the same buffer (2 drops/mL and 1 drop/mL, respectively). Anti-Aspergillus rabbit polyclonal antibody (Abcam, Cat.# ab20419) was also added (1/200) in the staining buffer for the conditions where *A. niger* conidia were inoculated. Fixed cells were incubated with 100 μL buffer containing the stains and Anti-Mucin 5AC and Anti-Aspergillus antibodies overnight at 4°C. The next day, fixed cells were washed 3 times with DPBS and secondary antibodies were applied. Goat anti-Mouse IgG antibody (1/250) conjugated with Alexa Fluor 546 (Thermo Fisher Scientific, USA, Cat.# A-11003,) directed against Anti-Mucin 5AC antibody and Goat anti-Rabbit IgG antibody (1/500) conjugated with Alexa Fluor 594 (Thermo Fisher Scientific, USA, Cat.# A-11012) directed against Anti-Aspergillus antibody were prepared in DPBS. Fixed cells were incubated with 100 μL buffer containing the secondary antibodies overnight at 4°C. The following day, fixed cells were once again washed 3 times with DPBS and the membranes from the inserts were carefully cut out with a sharp knife. The membranes were mounted on a glass slide using Fluoromount-G™ Mounting Medium (Thermo Fisher Scientific, USA) and imaged with a Zeiss Axio Observer Z1 fluorescence inverted microscope (Carl Zeiss AG, Germany).

### pH measurements and quantification of calcium, oxalic acid and lactate dehydrogenase (LDH)

pH measurements of the culture medium after 72h incubation were done directly after taking the samples out of the incubator using pH-indicator strips (Merck, Germany) for the Transwell® inserts samples, and CG8+ i-STAT cartridges for the BoC devices, as the pH change of the culture medium indicator was less visible. Free calcium (Calcium Colorimetric Assay, Sigma-Aldrich, Germany), free oxalic acid (Oxalic Acid Colorimetric Assay Kit), and LDH (ScienCell Research Laboratories, USA) were quantified in the culture medium using colorimetric assay kits following the manufacturer instructions.

### Statistical analyses

Statistical significance of the data from the confrontations on differentiated bronchial tissues in Transwell® inserts (3 replicates, n = 3) was tested with unpaired two-tailed Student t-tests in Microsoft® Excel (Version 16.37). The statistical significance threshold was set to 5%.

## Acknowledgments

We would like to thank Diego Gonzales and Ted Turlings for critical review of the paper, and Pulak Nath from the Materials Physics and Applications Division of the Los Alamos National Laboratory for providing the equipment and laboratory infrastructure for the BoC devices fabrication. BoC devices were developed under Defense Threat Reduction Agency (DTRA) interagency agreement CBMXCEL-XL1-2-0001.

## Data and materials availability

All data is available in the main text or the supplementary materials.

## References

1. Fisher MC, Gurr SJ, Cuomo CA, Blehert DS, Jin H, Stukenbrock EH, et al. Threats posed by the fungal kingdom to humans, wildlife, and agriculture. mBio. 2020;11(3).

2. Bongomin F, Gago S, Oladele RO, Denning DW. Global and multi-national prevalence of fungal diseases—estimate precision. 2017.

3. Ramirez-Ortiz ZG, Means TK. The role of dendritic cells in the innate recognition of pathogenic fungi (A. fumigatus, C. neoformans and C. albicans). 2012. p. 635–46.

4. Fisher MC, Hawkins NJ, Sanglard D, Gurr SJ. Worldwide emergence of resistance to antifungal drugs challenges human health and food security. 2018. p. 739–42.

5. Brown GD, Denning DW, Gow NAR, Levitz SM, Netea MG, White TC. Hidden killers: Human fungal infections. 2012. p. 165rv13–rv13.

6. Brunke S, Mogavero S, Kasper L, Hube B. Virulence factors in fungal pathogens of man. 2016. p. 89–95.

7. Pfaller MA, Diekema DJ. Rare and emerging opportunistic fungal pathogens: Concern for resistance beyond Candida albicans and Aspergillus fumigatus. 2004. p. 4419–31.

8. Kousha M, Tadi R, Soubani AO. Pulmonary aspergillosis: A clinical review. 2011. p. 156–74.

9. Paulussen C, Hallsworth JE, Álvarez-Pérez S, Nierman WC, Hamill PG, Blain D, et al. Ecology of aspergillosis: insights into the pathogenic potency of Aspergillus fumigatus and some other Aspergillus species. Microbial Biotechnology. 2017;10(2):296–322.

10. Kurrein F, Path FRC, Green GH, Rowles SL. Localized deposition of calcium oxalate around a pulmonary Aspergillus niger fungus ball. American Journal of Clinical Pathology. 1975;64(4):556–63.

11. Maeno T, Sasaki M, Shibue Y, Mimura K, Oka H. Calcium oxalate in the sputum may aid in the diagnosis of pulmonary aspergillosis: A report of two cases. Medical Mycology Case Reports. 2015;8:32–6.

12. Muntz FHA. Oxalate-producing pulmonary aspergillosis in an alpaca. Veterinary Pathology. 1999;36(6):631–2.

13. Oda M, Saraya T, Wakayama M, Shibuya K, Ogawa Y, Inui T, et al. Calcium oxalate crystal deposition in a patient with Aspergilloma due to Aspergillus niger. Journal of thoracic disease. 2013;5(4):E174–8.

14. Payne CL, Dark MJ, Conway JA, Farina LL. A retrospective study of the prevalence of calcium oxalate crystals in veterinary Aspergillus cases. Journal of Veterinary Diagnostic Investigation. 2017;29(1):51–8.

15. Yi Y, Cho SY, Lee DG, Jung JI, Park YJ, Lee KY. Invasive Pulmonary Aspergillosis Due to Aspergillus awamori: Role of Calcium Oxalate Crystal Precipitation Mimicking Mucormycosis. 2020. p. 409–11.

16. Cessna SG, Sears VE, Dickman MB, Low PS. Oxalic acid, a pathogenicity factor for Sclerotinia sclerotiorum, suppresses the oxidative burst of the host plant. Plant Cell. 2000;12(11):2191–9.

17. Schoonbeek H-j, Jacquat-Bovet A-C, Mascher F, Métraux J-P. Oxalate-degrading bacteria can protect Arabidopsis thaliana and crop plants against Botrytis cinerea. Molecular plant-microbe interactions. 2007;20(12):1535–44.

18. Lehner A, Meimoun P, Errakhi R, Madiona K, Barakate M, Bouteau F. Toxic and signalling effects of oxalic acid. Canadian Journal of Microbiology. 2008;3(September):746–8.

19. Dutton MV, Evans CS. Oxalate production by fungi: its role in pathogenicity and ecology in the soil environment. Canadian Journal of Microbiology. 1996;42(9):881–95.

20. Palmieri F, Estoppey A, House GL, Lohberger A, Bindschedler S, Chain PSG, et al. Oxalic acid, a molecule at the crossroads of bacterial-fungal interactions. In: Gadd GM, Sariaslani S, editors. 106: Academic Press; 2019. p. 49–77.

21. de Oliveira Ceita G, Macêdo JNA, Santos TB, Alemanno L, da Silva Gesteira A, Micheli F, et al. Involvement of calcium oxalate degradation during programmed cell death in Theobroma cacao tissues triggered by the hemibiotrophic fungus Moniliophthora perniciosa. Plant Science. 2007;173(2):106–17.

22. Graustein WC, Cromack K, Sollins P. Calcium oxalate: Occurrence in soils and effect on nutrient and geochemical cycles. Science. 1977;198(4323):1252–4.

23. Martin G, Guggiari M, Bravo D, Zopfi J, Cailleau G, Aragno M, et al. Fungi, bacteria and soil pH: The oxalate-carbonate pathway as a model for metabolic interaction. Environmental Microbiology. 2012;14(11):2960–70.

24. Hofmann BA, Bernasconi SM. Review of occurrences and carbon isotope geochemistry of oxalate minerals: implications for the origin and fate of oxalate in diagenetic and hydrothermal fluids. Chemical Geology. 1998;149(1-2):127–46.

25. Certini G, Corti G, Ugolini FC. Vertical trends of oxalate concentration in two soils under Abies alba from Tuscany (Italy). Journal of Plant Nutrition and Soil Science. 2000;163(2):173–7.

26. Braissant O, Cailleau G, Aragno M, Verrecchia EP. Biologically induced mineralization in the tree Milicia excelsa (Moraceae): its causes and consequences to the environment. Geobiology. 2004;2(1):59–66.

27. Rudnick MB, van Veen JA, de Boer W. Oxalic acid: A signal molecule for fungus-feeding bacteria of the genus Collimonas? Environmental Microbiology Reports. 2015;7(5):709–14.

28. Mela F, Fritsche K, De Boer W, Van Veen JA, De Graaff LH, Van Den Berg M, et al. Dual transcriptional profiling of a bacterial/fungal confrontation: Collimonas fungivorans versus Aspergillus niger. ISME Journal. 2011;5(9):1494–504.

29. Haq IU, Zwahlen RD, Yang P, van Elsas JD. The Response of Paraburkholderia terrae Strains to Two Soil Fungi and the Potential Role of Oxalate. Frontiers in Microbiology. 2018;9(989).

30. Dethlefsen L, McFall-Ngai M, Relman DA. An ecological and evolutionary perspective on human-microbe mutualism and disease. Nature. 2007;449(7164):811–8.

31. Schuster E, Dunn-Coleman N, Frisvad J, Van Dijck P. On the safety of Aspergillus niger - A review. 2002. p. 426–35.

32. Ruijter GJG, van de Vondervoort PJI, Visser J. Oxalic acid production by Aspergillus niger: an oxalate-non-producing mutant produces citric acid at pH 5 and in the presence of manganese. Microbiology. 1999;145(9):2569–76.

33. Cameselle C, Bohlmann JT, Núñez MJ, Lema JM. Oxalic acid production by Aspergillus niger. Bioprocess Engineering. 1998;19(4):247–52.

34. Person AK, Chudgar SM, Norton BL, Tong BC, Stout JE. Aspergillus niger: An unusual cause of invasive pulmonary aspergillosis. Journal of Medical Microbiology. 2010;59(7):834–8.

35. Plassard C, Fransson P. Regulation of low-molecular weight organic acid production in fungi. 2009. p. 30–9.

36. Show PL, Oladele KO, Siew QY, Aziz Zakry FA, Lan JCW, Ling TC. Overview of citric acid production from Aspergillus niger. 2015. p. 271–83.

37. Sturm EV, Frank-Kamenetskaya O, Vlasov D, Zelenskaya M, Sazanova K, Rusakov A, et al. Crystallization of calcium oxalate hydrates by interaction of calcite marble with fungus Aspergillus Niger. American Mineralogist. 2015;100(11-12):2559–65.

38. Karaffa L, Kubicek CP. Aspergillus niger citric acid accumulation: do we understand this well working black box? Appl Microbiol Biotechnol. 2003;61(3):189–96.

39. Kobayashi K, Hattori T, Honda Y, Kirimura K. Oxalic acid production by citric acid-producing Aspergillus niger overexpressing the oxaloacetate hydrolase gene oahA. J Ind Microbiol Biotechnol. 2014;41(5):749–56.

40. Fulcher ML, Gabriel S, Burns KA, Yankaskas JR, Randell SH. Well-differentiated human airway epithelial cell cultures. Methods in molecular medicine. 2005;107:183–206.

41. Kendall LV, Owiny JR, Dohm ED, Knapek KJ, Lee ES, Kopanke JH, et al. Replacement, Refinement, and Reduction in Animal Studies With Biohazardous Agents. ILAR journal. 2018;59(2):177–94.

42. Han Y, Joosten HJ, Niu W, Zhao Z, Mariano PS, McCalman M, et al. Oxaloacetate hydrolase, the C-C bond lyase of oxalate secreting fungi. Journal of Biological Chemistry. 2007;282(13):9581–90.

43. Abratt VR, Reid SJ. Oxalate-degrading bacteria of the human gut as probiotics in the management of kidney stone disease. 72: Academic Press; 2010. p. 63–87.

44. Hoppe B, Von Unruh G, Laube N, Hesse A, Sidhu H, editors. Oxalate degrading bacteria: New treatment option for patients with primary and secondary hyperoxaluria?2005/11//.

45. Enaud R, Prevel R, Ciarlo E, Beaufils F, Wieërs G, Guery B, et al. The Gut-Lung Axis in Health and Respiratory Diseases: A Place for Inter-Organ and Inter-Kingdom Crosstalks. 2020.

46. Mitchell AB, Oliver BGG, Glanville AR. Translational aspects of the human respiratory virome. 2016. p. 1458–64.

47. Turroni S, Vitali B, Bendazzoli C, Candela M, Gotti R, Federici F, et al. Oxalate consumption by lactobacilli: Evaluation of oxalyl-CoA decarboxylase and formyl-CoA transferase activity in Lactobacillus acidophilus. Journal of Applied Microbiology. 2007;103(5):1600–9.

48. Miller AW, Dearing D. The metabolic and ecological interactions of oxalate-degrading bacteria in the mammalian gut. 2013. p. 636–52.

49. Ticinesi A, Nouvenne A, Chiussi G, Castaldo G, Guerra A, Meschi T. Calcium oxalate nephrolithiasis and gut microbiota: Not just a gut-kidney axis. a nutritional perspective. 2020.

50. Suryavanshi MV, Bhute SS, Jadhav SD, Bhatia MS, Gune RP, Shouche YS. Hyperoxaluria leads to dysbiosis and drives selective enrichment of oxalate metabolizing bacterial species in recurrent kidney stone endures. Scientific Reports. 2016;6(1).

51. Pabuççuo□lu U. Aspects of oxalosis associated with aspergillosis in pathology specimens. Pathology Research and Practice. 2005;201(5):363–8.

52. Bertuzzi M, van Rhijn N, Krappmann S, Bowyer P, Bromley MJ, Bignell EM. On the lineage of Aspergillus fumigatus isolates in common laboratory use. Medical Mycology. 2020:myaa075-myaa.

53. Hsieh HL, Nath P, Huang JH. Multistep Fluidic Control Network toward the Automated Generation of Organ-on-a-Chip. ACS Biomaterials Science and Engineering. 2019;5(9):4852–60.

54. Lin CK, Hsiao YY, Nath P, Huang JH. Aerosol delivery into small anatomical airway model through spontaneous engineered breathing. Biomicrofluidics. 2019;13(4):044109-.

55. Arefin A, McCulloch Q, Martinez R, Martin SA, Singh R, Ishak OM, et al. Micromachining of Polyurethane Membranes for Tissue Engineering Applications. ACS Biomaterials Science and Engineering. 2018;4(10):3522–33.

56. Nelson KE, Weinel C, Paulsen IT, Dodson RJ, Hilbert H, Martins dos Santos VAP, et al. Complete genome sequence and comparative analysis of the metabolically versatile Pseudomonas putida KT2440. Environmental Microbiology. 2002;4(12):799–808.

57. Sentchilo V, Czechowska K, Pradervand N, Minoia M, Miyazaki R, Van Der Meer JR. Intracellular excision and reintegration dynamics of the ICEclc genomic island of Pseudomonas knackmussii sp. strain B13. Molecular Microbiology. 2009;72(5):1293–306.

58. Şahin N, Işik K, Tamer AÜ, Goodfellow M. Taxonomic position of ‘Pseudomonas oxalaticus’ strain Ox1(T) (DSM 110(T)) (Khambata and Bhat, 1953) and its description in the genus Ralstonia as Ralstonia oxalatica comb, nov. Systematic and Applied Microbiology. 2000;23(2):206–9.

59. Imeria Ferro K. The impact of oxalogenic plants on soil carbon dynamics - formation of a millennium carbon storage as calcium carbonate. Neuchâtel: University of Neuchâtel; 2012.

60. Reasoner DJ, Geldreich EE. A new medium for the enumeration and subculture of bacteria from potable water. Applied and Environmental Microbiology. 1985;49(1):1–7.

