## Supplementary Material for "Biocontrol of *Aspergillus niger* in 3D-lung cell tissues by oxalotrophic bacteria"

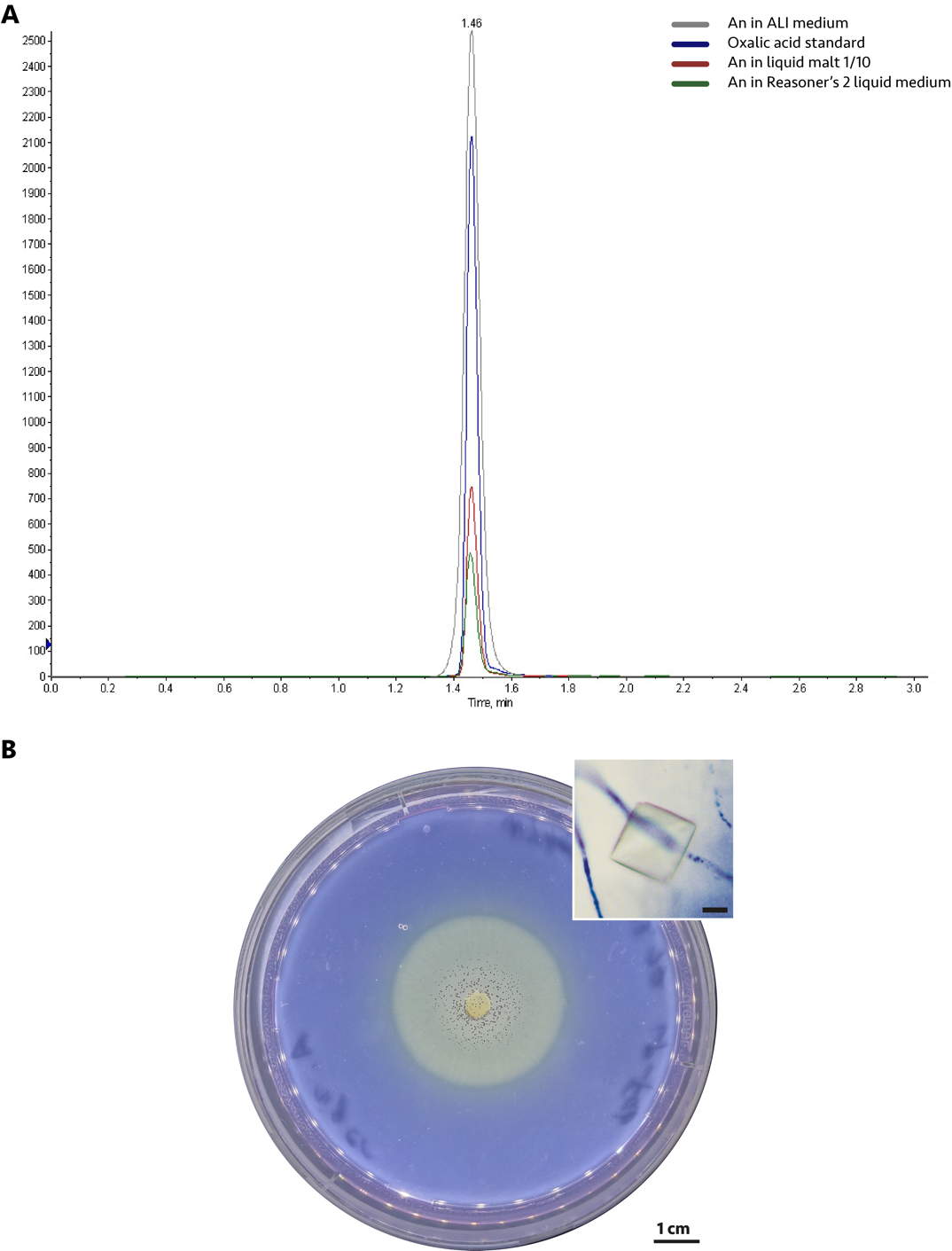

**Fig. S1.** **Detection of oxalic acid production by *Aspergillus niger* in different culture media.** (A) Ultra-High-Performance Liquid Chromatography (UHPLC) analysis confirmed the production of oxalic acid (blue chromatogram) by *A. niger* in Air-Liquid Interface (ALI, gray chromatogram), liquid malt 1/10 (red chromatogram) and liquid reasoner’s 2 media (green chromatogram). (B) Production of low molecular weight organic acids (LMWOA) by *A. niger* in a water yeast agar (WYA) plate supplemented with bromocresol purple as pH indicator. A yellow color indicates an acidic pH. The presence of characteristic bipyramidal shaped calcium oxalate crystals in the agar medium confirmed the production of oxalic acid by *A. niger* in this medium. Scale bar in the insert = 10 μm.

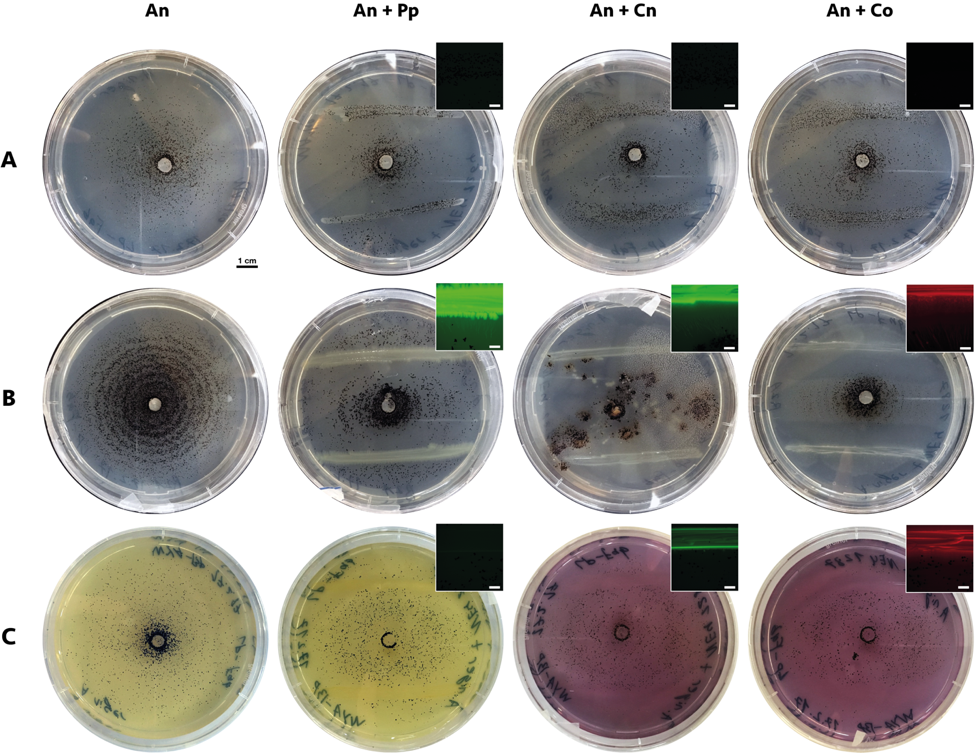
**Fig. S2.** **Interaction between *A. niger* (An), the non-oxalotrophic bacterium *P. putida* (Pp) and the oxalotrophic bacteria *C. necator* (Cn) and *C. oxalaticus* (Co) on different culture media.** Growth (and pH) inhibition depends on the nutrient medium. (A) In medium favoring the fungus (malt agar diluted 1/10 -MA 1/10-), the bacteria are all killed, regardless of their metabolism. Indeed, the lack of fluorescence (see inserts) confirms bacterial death. (B) In medium favoring the bacteria (R2A), oxalotrophic bacteria seem to control fungal growth, in contrast to the non-oxalotrophic strain. (C) In a poor-nutrient medium (WYA supplemented with Bromocresol Purple pH indicator), the oxalotrophic bacteria seem to control the LMWOA production by the fungus, as showed by the color reversal of the pH indicator (a yellow color indicates an acidic pH <6). As confirmed by the epifluorescence stereoscope observation, both oxalotrophic strains are alive, whereas the non-oxalotrophic one is dead. Scale bars in inserts: 2000µm.

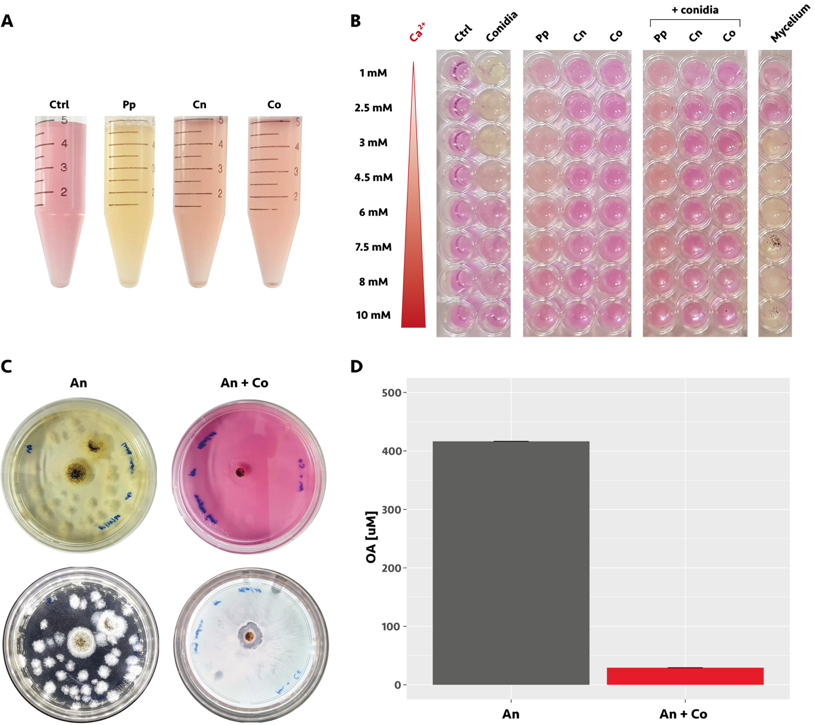

**Fig. S3.** **Growth tests and confrontation assay of *Aspergillus niger* and *C. oxalaticus* in Air-Liquid Interface (ALI) medium**. (A). The oxalotrophic bacteria (*Cupriavidus necator* –Cn- and *C. oxalaticus* –Co-) developed in ALI medium but without inducing a change in pH in contrast to non-oxalotrophic bacteria (*Pseudomonas putida* –Pp-). pH indicator= phenol red. (B). Development of the same bacterial species indicated in A, as well as *A. niger* inoculated as conidia or mycelium on ALI medium with increasing concentrations of calcium. Calcium concentration and the mode of inoculation (fungal conidia versus mycelium) has an effect on fungal development and control of fungal growth by bacteria. While conidia germination was inhibited at 6 mM Ca^2+^ and above, mycelial growth was stimulated from 3-10 mM, but inhibited at the lower Ca^2+^ concentrations. Growth of *P. putida* was also affected by Ca^2+^ (inhibition above 8 mM), but increasing Ca^2+^ concentrations did not affect growth of the oxalotrophic bacteria. When conidia were co-inoculated with the three bacterial species indicated above, all the bacteria appear to partially inhibit germination, but the effect on the stabilization of the medium pH was only observed with the two oxalotrophic bacteria. (C) The top row shows the difference in pH (yellow for acidic pH and pink for neutral) for both treatments: fungus alone (An) and fungus in co-culture with the bacterium (An+Co). The bottom row shows images of same plates on the top, but in this case, a light source was used from below the plate to better visualize the growth of the fungus in both conditions. (D) Oxalic acid concentration decreased by close to 90% in presence of the bacterium (An+Co) as compared to the control with the fungus alone (An).

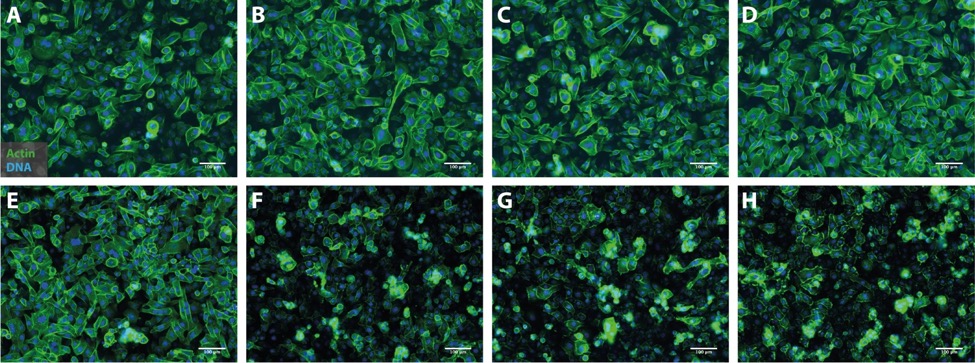
**Fig. S4.** **Effect of increasing *A. niger* conidial load on the morphology of bronchial epithelial cells.** (A) Control cells. (B) 10 conidia. (C) 50 conidia. (D) 100 conidia. (E) 500 conidia. (F) 1000 conidia. (G) 5000 conidia. (H) 10’000 conidia. After 24h incubation, an increase in cell damage with increasing conidial load was observed. Damage was visible from a conidial load of 500. Damaged cells began to shrink, and actin got more agglomerated, compared to the cells-only control. From a conidial load of 1000 and on, fungal growth has an adverse effect on tissue integrity. Culture medium volume was 200 μl/well. Scale bars = 100 μm.

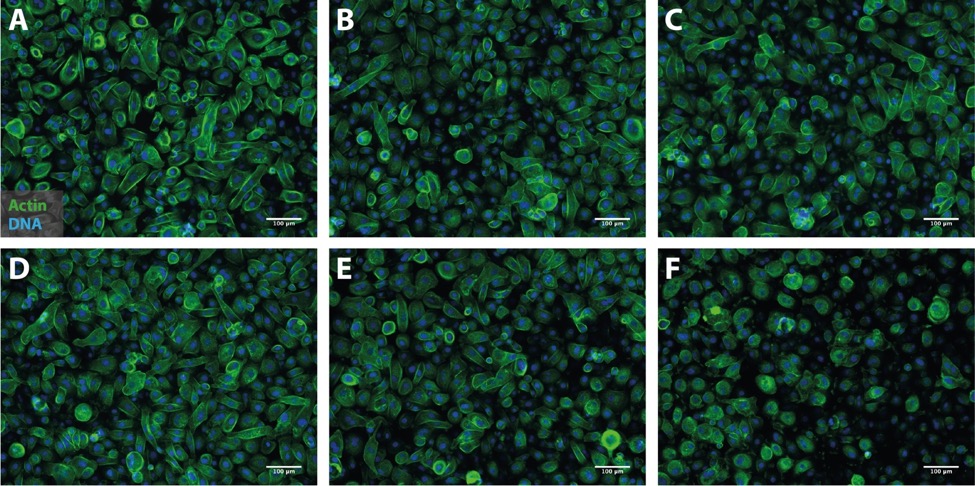

**Fig. S5.** **Effect of increasing cell load of *C. oxalaticus* on the morphology of bronchial epithelial cells.** (A) Control cells. (B) 10 bacterial cells. (C) 50 bacterial cells. (D) 100 bacterial cells. (E) 500 bacterial cells. (F) 1000 bacterial cells. After 24h incubation, an increase in cell damage with increasing bacterial cell load was observed. As few as 10 bacterial cells have already an impact on cell morphology. Indeed, cells became rounder, and actin got more agglomerated, compared to the cells-only control. However, the cytopathic effect observed in the presence of *A. niger* was less pronounced (Figure S4). Culture medium volume was 200 μl/well. Scale bars = 100 μm.

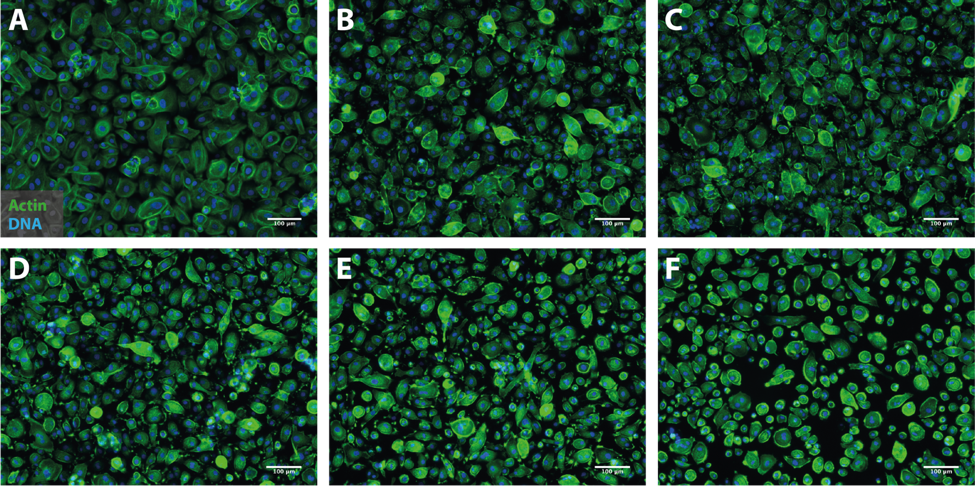
 **Fig. S6.** **Effect of increasing cell load of *P. putida* KT2440 on the morphology of bronchial epithelial cells.** (A) Control cells. (B) 10 bacterial cells. (C) 50 bacterial cells. (D) 100 bacterial cells. (E) 500 bacterial cells. (F) 1000 bacterial cells. After 24h incubation, an increase in cell damage with increasing bacterial cell load was observed. As few as 10 bacterial cells have already a strong impact on cell morphology. Indeed, cells became rounder, and actin got more agglomerated, compared to the cells-only control. Culture medium volume was 200 μl/well. Scale bars = 100 μm.

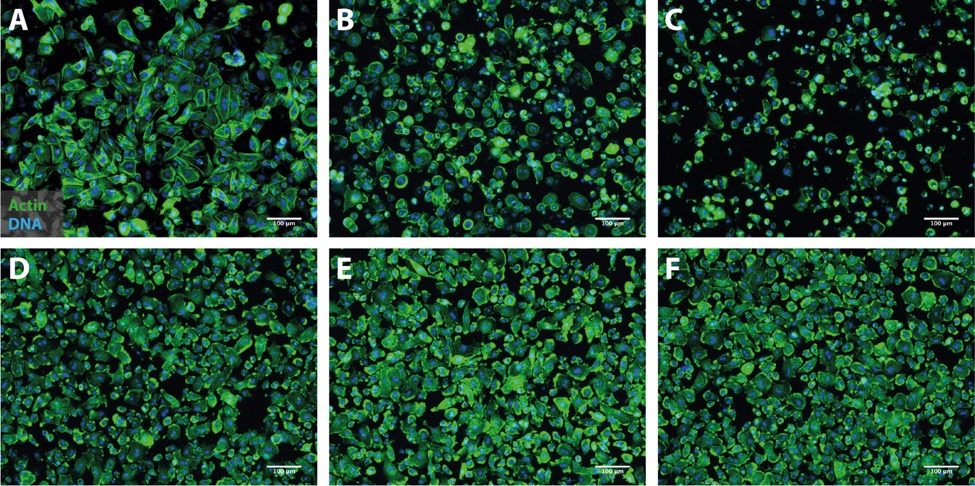
**Fig. S7.** **Effect of co-culturing *C. oxalaticus* with *A. niger* on cell morphology in submerged undifferentiated bronchial epithelium.** (A) Cell control. (B) 10 *A. niger* conidia. (C) 500 *A. niger* conidia. (D) 10 *C. oxalaticus* cells. (E) 10 *A. niger* conidia **+** 10 *C. oxalaticus* cells. (F) 500 *A. niger* conidia **+** 10 *C. oxalaticus* cells. After 72h incubation, the damage and cytopathic effect of *A. niger* conidia was clearly visible with as few as 10 conidia per well (200 μl) (B). The cells appear even more damaged with a conidial load of 500 (C). Ten *C. oxalaticus* cells also changed the morphology of the epithelial cells, but no cytopathic effect was observed (D). With the co-inoculation of as few as 10 *C. oxalaticus* cells (E and F), the morphology of bronchial cells infected with *A. niger* was similar the morphology of bacteria-only control. Culture medium volume was 200 μl/well. Scale bars = 100 μm.

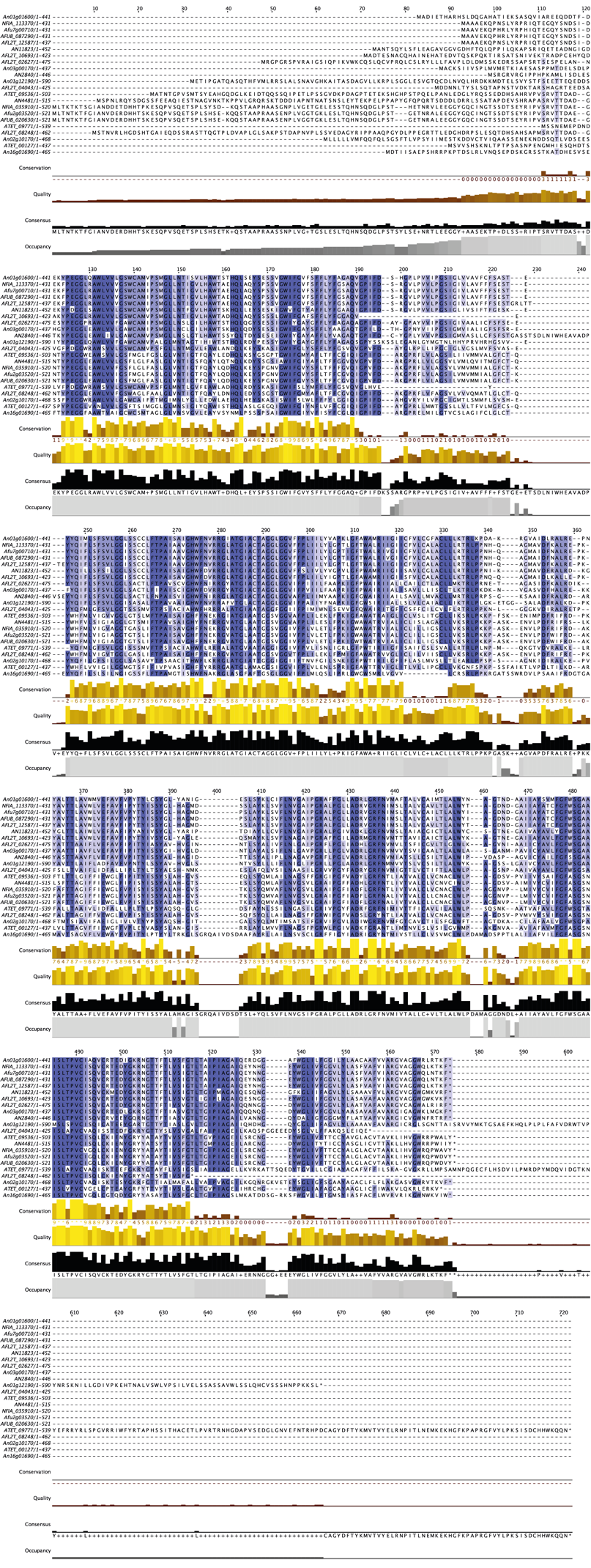

**Fig. S8.** **Genomic screening of the oxalate/formate antiporter in other *Aspergillus* spp.** Multiple sequence alignment of the protein sequences orthologous to the oxalate/formate antiporter of *A. niger* CBS 513.88 (GenBank accession number XP_001388599.2) revealed they were conserved to a lesser extent, as indicated by an intense purple color of the amino acids. Multiple sequence alignments were performed using the MUSCLE protein alignment algorithm in Jalview (version 2.11.1.2). An = *A. niger* CBS 513.88, NFIA = *Neosartorya fisheri* NRRL 181 (formerly *A. fisheri*), Afu = *A. fumigatus* Af293, AFUB = *A. fumigatus* A1163, AFL2T = *A. flavus* NRRL 3357, AN = *A. nidulans* FGSC A4, ATET = *A. terreus* NIH2624.
